# The pregnant myometrium is epigenetically activated at contractility-driving gene loci prior to the onset of labor in mice

**DOI:** 10.1101/2020.03.19.998740

**Authors:** Virlana M. Shchuka, Luis E. Abatti, Huayun Hou, Anna Dorogin, Michael D. Wilson, Oksana Shynlova, Jennifer A. Mitchell

**Affiliations:** Department of Cell and Systems Biology, University of Toronto, Toronto, ON, M5S 3G5, Canada; Department of Molecular Genetics, University of Toronto, Toronto, ON, M5S 1A8, Canada; Genetics and Genome Biology Program, SickKids Research Institute, Toronto, ON, M5G 0A4, Canada; Lunenfeld Tanenbaum Research Institute, Sinai Health System, Toronto, ON, M5G 1X5, Canada; Department of Obstetrics & Gynaecology, University of Toronto, ON, M5G 1E2, Canada

**Author notes:** These authors contributed equally to the work (Virlana M. Shchuka, Luis E. Abatti). Department of Cell and Systems Biology, University of Toronto, Toronto, Canada.

## Abstract

During gestation, uterine smooth muscle cells transition from a state of quiescence to one of contractility, but the molecular mechanisms underlying this transition at a genomic level are not well-known. To better understand these events, we evaluated the epigenetic landscape of the mouse myometrium during pregnancy, labor and post-partum. We established gestational timepoint-specific enrichment profiles involving histone H3K27 acetylation (H3K27ac), H3K4 tri-methylation (H3K4me3), and RNA polymerase II (RNAPII) occupancy by chromatin immunoprecipitation sequencing (ChIP-seq), as well as gene expression profiles by total RNA-sequencing (RNA-seq). Our findings reveal that 533 genes, including known contractility-driving genes (*Gja1, Fos, Oxtr, Ptgs2*), are upregulated during active labor due to an increase in transcription at gene bodies. Their promoters and putative intergenic enhancers, however, are epigenetically activated by H3K27ac as early as day 15, four days prior to the onset of active labor on day 19. In fact, we find that the majority of genome-wide H3K27ac or H3K4me3 peaks identified during active labor are present in the myometrium on day 15. Despite the early presence of H3K27ac at labor-associated genes, both an increase in non-coding enhancer RNA (eRNA) production, and in recruitment of RNAPII to corresponding genes occur during active labor, at labor upregulated gene loci. Our findings indicate that epigenetic activation of the myometrial genome precedes active labor by at least four days in the mouse model, suggesting the myometrium is poised for rapid activation of contraction-associated genes in order to exit the state of quiescence.

## Introduction

Over the course of gestation, the myometrium transitions from a state of quiescence during pregnancy to one of contractile activity during labor in response to both hormonal and mechanical signals. Concomitant changes in gene expression that accompany this transition are thought to be a driving force for the initiation of labor [1,2]; however, little is known about the molecular mechanisms underlying these changes. Across developmental contexts, the chromatin landscape is thought to maintain a cell’s identity, with dynamic chromatin state changes differentiating various cell types from one another as well as the same cell type under different conditions [3]. Across cell types, transcription start sites (TSS) of actively transcribed genes are marked by histone H3 tri-methylation of lysine 4 (H3K4me3) and acetylation on lysine 27 (H3K27ac) and increased gene expression levels are correlated with the presence of both markers at gene TSS [4–8]. Additionally, transcriptional enhancers, which can be located at kilobase-to megabase-sized distances from the genes they regulate, contain a prominent signature consisting of H3K27ac [3,4,9–11] and non-coding enhancer RNAs (eRNAs), both of which can be used as a means of identifying regions with tissue-specific enhancer activity [12–14]. Finally, the presence of histone modifications typically associated with active genes and the subsequent recruitment of RNA polymerase II (RNAPII) to gene promoters allow for transcription initiation and transition to elongation, thereby upregulating gene expression [15]. Where, how, and at what point these events occur in the myometrial genome during gestation are the inquiries guiding this study.

We know that uterine contractions are enabled when myometrial muscle cells act *en masse* to generate a series of synchronous movements, actions that require the coupling of cells through the presence of intercellular bridges, or gap junctions. Among the proteins mediating junction formation as term approaches, gap junction alpha 1 (GJA1, also known as CX43) is most prominently upregulated [16]. Selective reduction of GJA1 production in the uterine smooth muscle cells of two different mouse models has been shown to significantly prolong the quiescent state during pregnancy and thereby delay the onset of labor [17,18]. Reporter expression downstream of a synthetic *Gja1* promoter is increased by co-expression of constructs encoding members of the activator protein 1 (AP-1) transcription factor FOS and JUN sub-families [19–21]. Furthermore, increased levels of FOS and FOSL2 in particular within the nuclei of myometrial cells during labor raises the possibility that the FOS:JUN family acts to transcriptionally activate *Gja1* to initiate labor onset [22,23]. Several JUN sub-family members are present in the myometrium throughout gestation; however they display a more limited ability to act as activators of *Gja1* promoter-driven transcription. It is therefore likely that JUN proteins may have a role in maintaining myometrial gene expression during pregnancy, but require heterodimerization with a FOS sub-family partner to activate genes required for the onset of labor.

Despite extensive *in vitro* studies correlating FOS:JUN activity with *Gja1* promoter activation and consequent labor initiation, little is known about the active chromatin landscape on a genome-wide scale in the myometrium as uterine smooth muscle cells exit the pregnant and enter the laboring state. We address this gap in the literature by investigating the epigenetic and transcriptomic changes that take place in the nucleus during this cellular transition. Using total RNA-sequencing (RNA-seq) methods, we observed an increase in primary transcript levels for the majority of genes that display increased expression during labor, suggesting that the initiation of contractility involves substantial modulation of gene transcription. Despite these temporally-dependent differences in transcription output, the myometrial genome does not undergo a corresponding acquisition of euchromatin-associated histone marks. Instead, we determined that H3K27ac and H3K4me3 modifications are established at labor-upregulated gene promoters during the uterine quiescent stage, several days prior to the onset of labor. Although gene promoters are pre-marked with these histone modifications, we identified increased RNAPII enrichment at promoters and across gene bodies, and increased expression of enhancer RNAs (eRNAs) in non-coding regions surrounding labor-associated genes during active labor. Furthermore, we found that intergenic regions exhibiting H3K27ac peaks and labor-upregulated eRNA expression displayed an enrichment of AP-1 transcription factor motifs, thereby implicating FOS:JUN heterodimers in the distal regulation of gene transcription changes at labor onset. These observations collectively suggest that the murine myometrium undergoes a cascade of epigenetic events that begins well in advance, and continues to the commencement, of labor at term.

## Results

### Up-regulation of labor-associated genes involves a transcriptional mechanism

To establish a comprehensive profile of pregnant and laboring myometrial transcriptomes, we conducted total strand-specific RNA-sequencing (RNA-seq) on RNA isolated from the myometrium of pregnant C57BL/6 mice at gestational day (d)15 or day (d)19 while in active labor (n=5 each, Fig 1A). Based on the RNA-seq data, we observed clustering of the same samples within each timepoint of collection, as expected (S1 Fig). Differential gene expression analysis based on exon read counts (S1 Table) revealed that a total of 956 genes showed gestational timepoint-varying expression levels (Fig 1B, fold change cut-off of 4, P<0.01), with hierarchical clustering analysis of these genes revealing similar expression trends in mice of the same gestational age (Fig 1C). 578 genes exhibited a significant increase in expression during active labor (d19) compared to d15. Apart from up-regulation of *Fos* (Fig 1D), these genes included (but were not limited to) prominent labor-associated players *Gja1, Ptgs2*, and *Oxtr*, as well as matrix metalloproteinases (*Mmp7, Mmp11, and Mmp12*); signaling proteins (*Cxcl1, Cxcl5*); and adhesion molecules and proteins (*Vcam1, Thbs1, Ceacam1*) known to exhibit elevated levels at term. Conversely, 378 genes were found to be significantly downregulated during active labor compared to d15, including (but not limited to) proteins responsible for cell-extracellular matrix interactions (*Col4a6, Col11a1, Col13a1, Col15a1, Col26a1, Spock2)*, proteins involved in calcium signaling (*Mchr1, Calml3, Calb2)*, proteins regulating myometrium response to low oxygen tension *(Hif3a)* and resistance to oxidative stress *(Akr1b7)*, and voltage-dependent calcium, potassium, and water channels *(Cacna1e, Kcng1*, and *Aqp8*, respectively).

**Figure 1.**
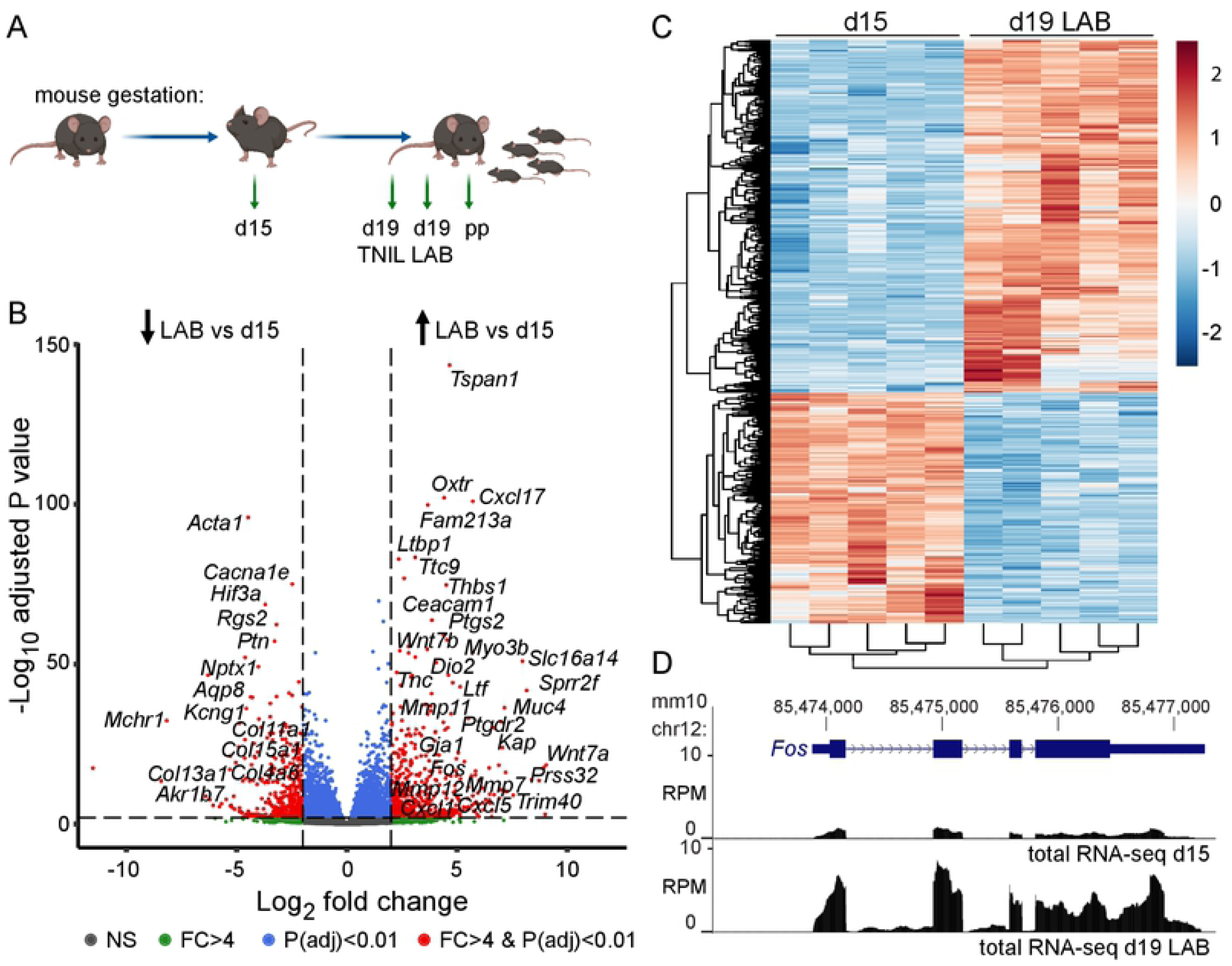
Quiescent and term laboring myometrial transcriptomes exhibit differential expression profiles. **(A)** Gestational schematic outlining days and timepoints at which myometrial tissues were collected for transcriptome- or genome-wide sequencing analyses. Collection days include gestational day 15 (d15), day 19 term-not-in-labor (d19 TNIL), day 19 active labor (d19 LAB) and postpartum (pp) (n=5 per gestational day). **(B)** RNA-seq volcano plot highlighting transcriptional status of genes exhibiting differential expression levels between d15 and d19 LAB myometrial tissues **(C)** Hierarchical clustering of gene groups based on RNA expression changes between d15 and d19 LAB samples. **(D)** Total RNA-seq reads (reads per million, RPM) at the labor-associated *Fos* gene locus for d15 and d19 LAB samples mapped to the mm10 mouse genome assembly.

Differential exonic RNA profiles, however, do not in and of themselves reflect a regulatory mechanism change at the level of transcription for those genes. Mediation of gene regulation can take place at multiple stages within a gene’s expression pathway prior to formation of a final translated and functional protein product. Apart from varying of levels of primary transcript generation in the nucleus, these mechanisms can include nuclear retention of processed mRNA [24,25], alternative splicing [26,27], and increased mRNA stability [28,29]. Furthermore, multiple gene activation and repression mechanisms can compete to determine the expression outcome of a particular gene, as demonstrated by select biological contexts in which there is an imperfect correlation between the levels of a gene’s transcribed and its translated products [30].

Since sequencing of total RNA provides a transcriptome-wide profile of reads spanning both exons and introns, both spliced and unspliced RNA species can be detected [31,32]. If critical labor-driving genes were significantly upregulated at the level of active transcription, we would expect to identify a substantial increase in reads corresponding to gene introns as well as those corresponding to their exons. Conversely, if activated genes were regulated exclusively or predominantly by post-transcriptional mechanisms such as RNA stability, we would expect that those genes’ corresponding intron reads would remain relatively constant throughout gestation. Given that a prior study has posited that mRNA stability may act as a critical player in regulation of *Gja1* in particular [33], we sought to definitively ascertain whether increased contractility-associated gene expression during labor involved any regulatory input from transcriptional mechanisms. Upon inspection of the *Fos* locus, we noted a striking labor-specific intronic RNA enrichment profile (Fig 2A). Subsequently, we conducted a genome-wide intron reads-concentrated gene expression analysis (iRNA-seq)[31], which uncovered multiple genes that displayed increased primary transcript generation at term (LAB) relative to day 15 (Fig 2B, S2 Table). In fact, the majority of genes (55%) up-regulated at labor on the basis of increased exon read accumulation also displayed a significant increase in intron reads (Fig 2C). Using exon-intron junction-spanning primers, we confirmed a significant increase in primary transcript levels of well-known labor-associated genes at d19 (LAB) relative to both day 15 and d19 term-not-in-labor (TNIL, Fig 2D). These results demonstrate that genes promoting the contractile state in the myometrium at term act due to a rapid gestational timepoint-specific increase in primary transcript levels, which suggests substantial transcriptional activity occurs in myometrial nuclei during labor.

**Figure 2.**
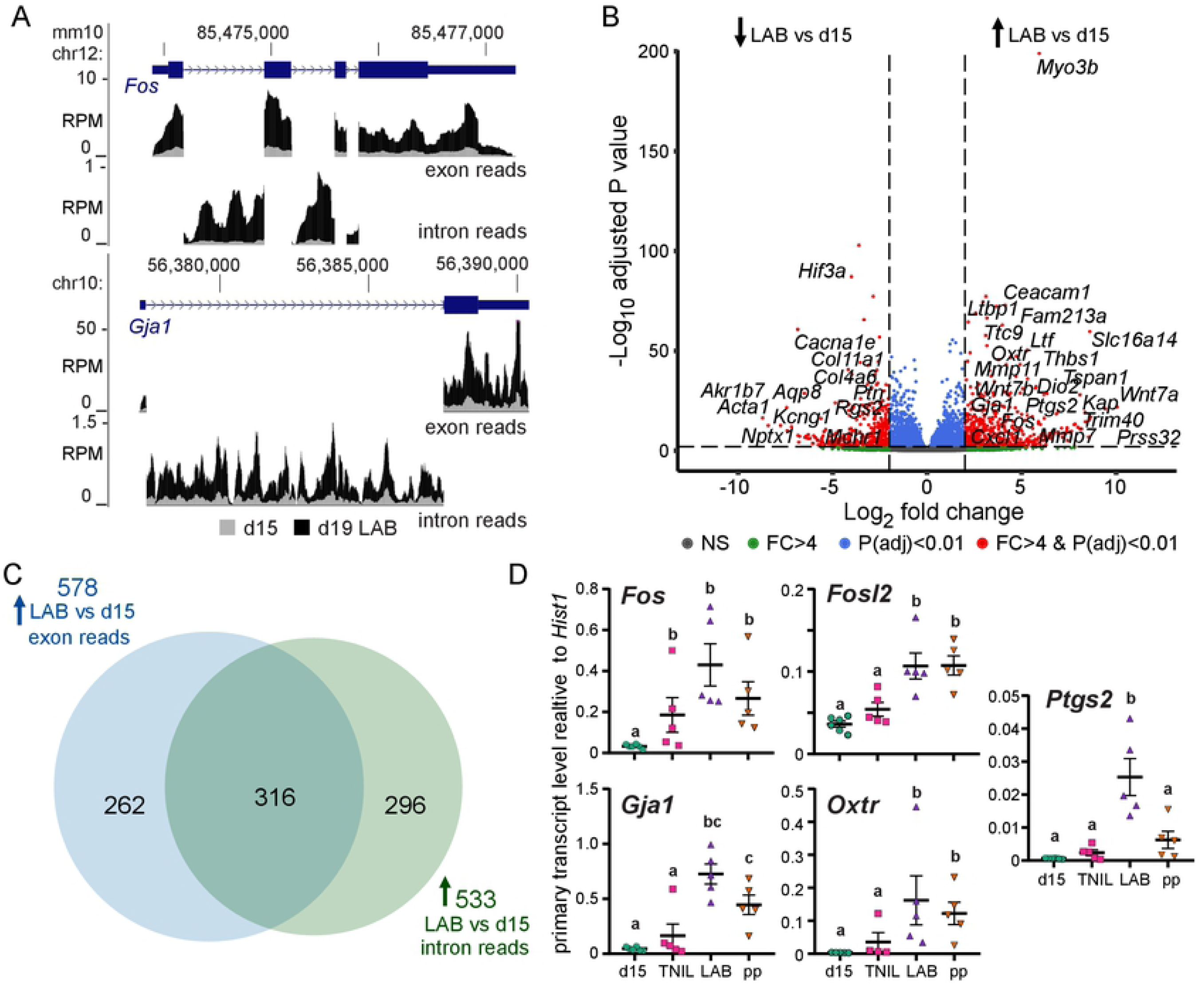
Up-regulation of labor-associated genes involves an increase in their primary transcript levels. **(A)** Exon- and intron-specific RNA-seq reads (reads per million, RPM) at the labor-associated *Fos* gene locus from d15 and d19 active laboring samples (d19 LAB), mapped to the mm10 mouse genome assembly. **(B)** Intron RNA-seq (iRNA-seq) volcano plot highlighting expression changes based on intron reads between day 15 (d15) and d19 active laboring (LAB) samples. **(C)** Venn diagram displaying the number of genes significantly up-regulated during labor relative to day 15 that show increased enrichment in exon and intron reads (region of overlap) and increased enrichment in either exon or intron reads (regions of non-overlap). **(D)** Confirmation of laboring timepoint-specific up-regulation of primary transcript expression of contractility-promoting genes by RT-qPCR. Groups labelled with different letters show significant difference, with p<0.05.

### Histone marks associated with gene activation are enriched at labor-associated gene promoters well in advance of labor onset

Having determined that the majority of genes exhibiting increased expression levels during labor were associated with increased primary transcript generation, we next investigated the active chromatin landscape surrounding these genes. After optimizing the protocol for myometrial tissue and confirming target enrichment at control regions (S2 Fig), we conducted chromatin immunoprecipitation and sequencing (ChIP-seq). We targeted H3K4me3 and H3K27ac enrichment events on a genome-wide scale at day 15, day 19 (TNIL), day 19 (LAB) and post-partum (pp), and found gestational timepoint samples replicate were highly correlated, as expected (R^2^ > 0.79, S3 Table). When we compared the histone profiles to the RNA-seq data, we observed that both H3K4me3 and H3K27ac are enriched at the promoters of highly expressed genes (S3 Fig). An initial examination of the *Fos* locus unexpectedly revealed an enrichment of both active chromatin markers at the *Fos* promoter across all four timepoints (Fig 3A). We next sought to establish the active histone profile across all gene promoters at the designated gestational stages (S4/S5 Tables). Even more surprisingly, we found a similar genome-wide active histone marker enrichment pattern at gene promoters (+/-2kb of TSS) in all four timepoints (Fig 3B). When we investigated the levels of these modifications at promoters of only those genes that displayed a significant increase in transcription (based on an increase in intron reads) during active labor, we observed only a moderate increase in accumulation of both markers from day 15 to term (Figs 3C and D). Our analyses therefore indicate that H3K27ac and H3K4me3 enrichment pre-marks the promoters of labor-upregulated genes as early as day 15 of gestation.

**Figure 3.**
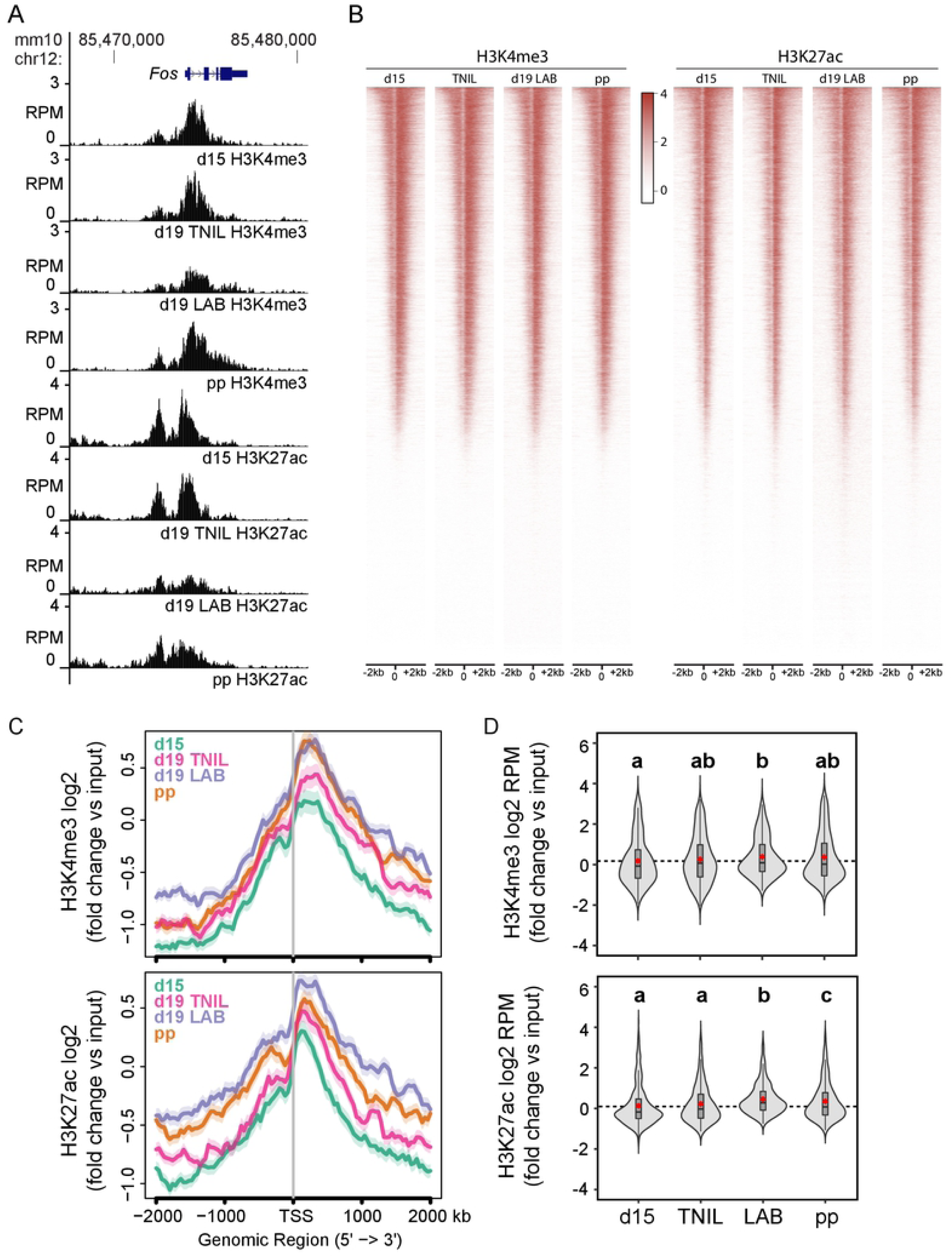
Gene expression activation-associated histone marks are enriched at promoters of contractility-driving genes in advance of labor onset. **(A)** Anti-H3K4me3 and anti-H3K27ac ChIP-seq reads (reads per million, RPM) mapped at promoter of labor-associated gene *Fos* in d15, d19 term-not-in-labor (TNIL), d19 active laboring (LAB), and postpartum (pp) samples. **(B)** Genome-wide enrichment of H3K4me3 and H3K27ac at gene promoters. Signal +/-2kb of TSS is displayed, with genes ordered at each indicated timepoint according to decreasing enrichment (red -> white) profile in d15 samples. **(C)** Plots exhibiting log_2_-fold H3K4me3 or H3K27ac signal, normalized to input, at promoters of genes whose expression is enriched in laboring samples relative to d15 samples based on intron reads. **(D)** Violin plots displaying log_2_-fold H3K4me3 or H3K27ac signal (reads per million, RPM), normalized to input, at promoters of genes whose intron read-based expression is enriched in laboring samples relative to d15 samples. Groups labelled with different letters show significant difference, with p<0.05.

### Loci of labor-associated genes acquire RNAPII gene body occupancy and eRNA enrichment events closer to term

Given that labor-associated gene loci exhibited strikingly similar active histone marker enrichment across all four gestational timepoints, we next sought to establish whether concomitant binding of RNAPII, an event required for transcription initiation, occurred at contractility-driving gene promoters and bodies as early in the gestational timecourse. We conducted ChIP-seq to identify RNAPII enrichment events (targeting its serine 5 phosphorylated RPB1 subunit) at gestational day 15 and d19 (LAB), and identified RNAPII-enriched broad peaks (S6 Table). Again, our correlation analysis confirmed the clustering of gestational timepoint replicates (S4 Fig), while an examination of RNAPII enrichment values alongside our RNA-seq data revealed that RNAPII has a more prominent binding profile at highly expressed genes (S5 Fig; S7 Table). Contrary to our activating histone mark analyses, we observed substantial differential RNAPII binding profiles at either gestational stage. We found that promoter and gene body polymerase occupancy at genes with significantly higher expression levels during labor was significantly higher at term (d19 LAB) relative to d15 (Figs 4A/B; S5 Fig). This result correlated with our observations of these genes’ gestational timepoint-specific primary transcript levels, and further supported the notion that transcriptional mechanisms underlie the rapid and prompt gene expression up-regulation events that underlie the myometrial state transition toward contractility.

**Figure 4.**
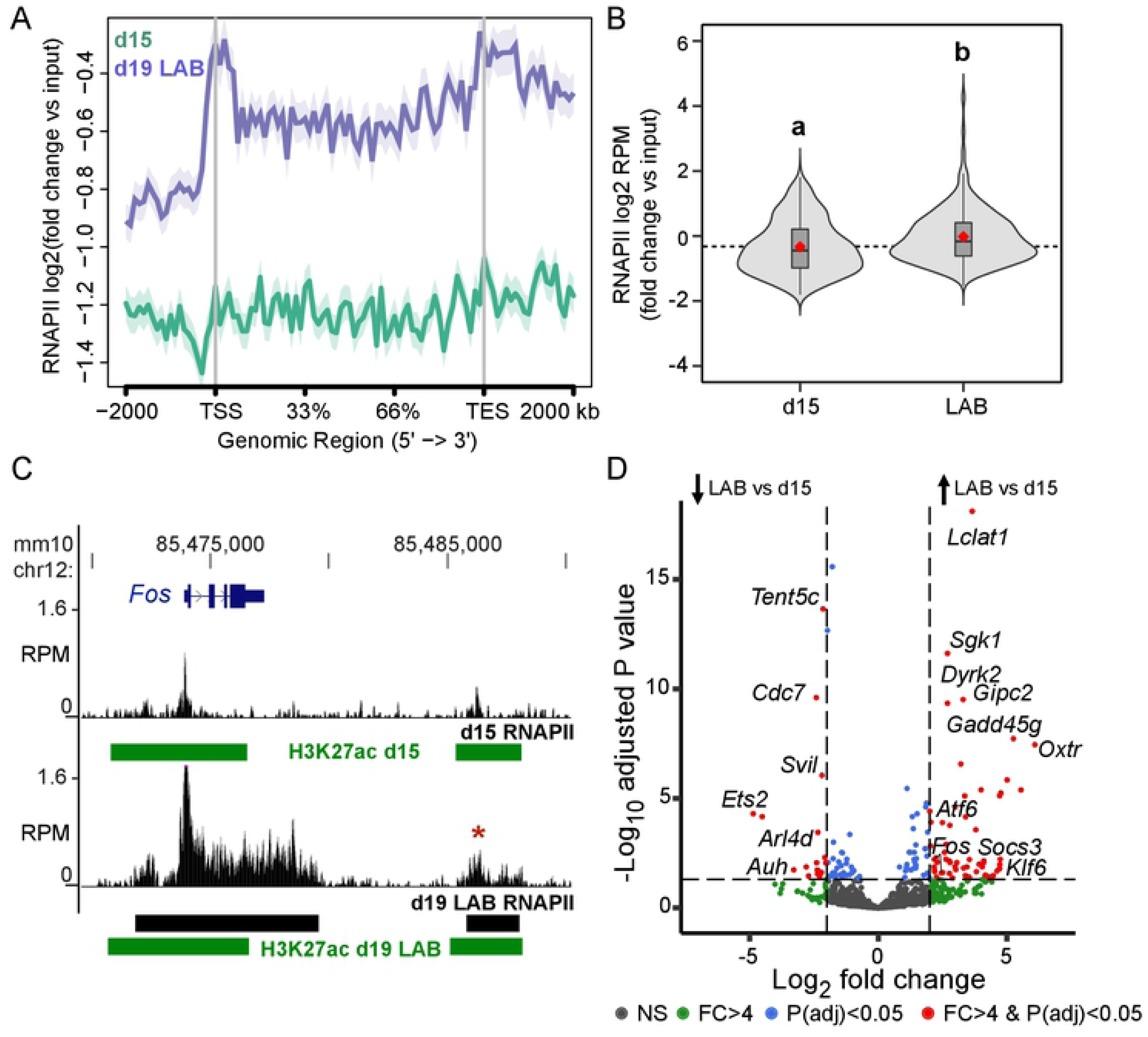
Active labor is associated with recruitment of RNAPII to labor-driving genes and eRNA expression at gene-adjacent intergenic regions with H3K27ac peaks. **(A)** Metagene plot exhibiting log_2_-fold RNAPII signal, normalized to input, at bodies of genes whose intron read-based expression levels are enriched in d19 laboring samples (d19 LAB) relative to d15 samples. **(B)** Violin plots displaying log_2_-fold RNAPII signal (reads per million, RPM), normalized to input, at promoters of genes whose intron read-based expression levels are enriched in laboring samples (LAB) relative to d15 samples. Groups labelled with different letters show significant difference, with p<0.05. **(C)** Anti-RNAPII ChIP-seq reads (reads per million, RPM) mapped at the *Fos* locus in d15 or d19 (LAB) samples. Regions containing H3K27ac peaks indicated in d15 and d19 (LAB) samples. Labor-up-regulated eRNA indicated at RNAPII-associated region downstream of gene. **(D)** Enhancer RNA (eRNA) volcano plot highlighting intergenic H3K27ac peaks in genomic regions that exhibit significant differences in eRNA levels between d15 and d19 laboring samples.

Among the genes significantly upregulated during labor, *Fos* exhibited increased RNAPII occupancy across its gene body, relative to day 15, as we expected. When we further expanded our view outside the gene body, we observed a genomic region that does not encode a gene, but includes both an RNAPII peak (* in Fig 4C) as well as an intergenic H3K27ac enrichment event 12kb downstream of the gene. As was the case with the H3K27ac signal at the *Fos* promoter, an intergenic H3K27ac peak was identified in both day 15 as well as d19 labor samples; however, RNAPII association at this region was more pronounced, and only identified as a peak in the labor context (Fig 4C). Since intergenic regions containing RNAPII and H3K27ac peaks have been noted to occur at active enhancer regions [6,12,34], we examined other intergenic regions of interest on a genome-wide scale. We found that, although we focused our earlier analyses on H3K27ac signal enrichment at promoter regions, 43% of the identified H3K27ac peaks in the laboring samples are located at distances greater than 2 kb from a gene TSS (Fig 4D, S8 Table). As was the case with H3K27ac peaks at gene promoters, intergenic H3K27ac peaks were mostly invariant across our tested gestational timepoints, with only 11/5041 (0.2%) of the intergenic H3K27ac peaks displaying a significant increase in H3K27ac signal enrichment in labor samples compared to day 15 samples (S9 Table). However, despite the presence of H3K27ac enrichment at intergenic regions across all four tested timepoints, many gene loci contain H3K27ac-modified regions with associated transcribed eRNAs (S10 Table). Furthermore, we observed that several of these regions contain H3K27ac peaks and display a significant increase in eRNA expression levels during labor (Fig 4D). Therefore, although most of H3K27ac peaks across the myometrial genome are present in both day 15 and at term, several regions containing those peaks transcribe significantly higher amounts of eRNA at term.

To identify the transcription factor motifs that could underlie the changes in gene expression that occur during labor, we conducted motif enrichment analyses using HOMER [35]. We uncovered a significant enrichment of AP-1 motifs (TGACTCA) in labor-associated intergenic regions (S6 Fig), implicating this family of transcription factors in a modulatory role with regards to enhancer activity at labor onset. We made a similar observation at promoters of genes displaying increased primary transcript abundance during labor (S7 Fig). Additionally, apart from AP-1 motifs, the promoters of these genes are enriched in several other motifs: TCF3 (E2A), an E-protein transcription factor that has been shown to assist co-activator proteins in the induction of gene transcription in other cell contexts [36]; CTCF, a zinc finger protein prominently known to bind promoter and enhancer regulatory elements [37]; and RELA (NFkB-p65), a member of the labor-associated NFkB-p65/IL-6 inflammatory pathway [38,39] that has also been affiliated with inducing transcription at the *Oxtr* promoter [40]. Taken together, these results lead us to propose that the controlled expression of labor-associated genes is driven by transcriptional regulation mechanisms, despite the apparent epigenetic activation of labor-associated loci well in advance of labor onset.

## Discussion

Based on our data, we propose a biological model wherein the myometrium’s preparation for labor at a genomic level begins well in advance of term (Fig 5). We propose that the presence of H3K27ac and H3K4me3 marks at labor-associated gene promoters in mouse myometrium renders them open and accessible as early as gestational day 15 (approximately three quarters of the way through the timecourse of mouse pregnancy), even though the expression of these genes is low at this gestational stage. The onset of labor, however, coincides with increased eRNA transcription within labor-associated gene loci at non-coding regions that are enriched for AP-1 sequence motifs and contain prominent H3K27ac peaks. Furthermore, the promoters of genes with increased expression during labor also contain an enrichment of these motifs, suggesting that the latter may allow for binding of AP-1 factors as well as phosphorylated RELA to both promoters and distal elements. We argue that these regulatory mechanisms enable gestational timepoint-specific recruitment of RNAPII and consequent primary transcript production, events that form the basis of the myometrial organ’s transition from a quiescent to a contractile state.

**Figure 5.**
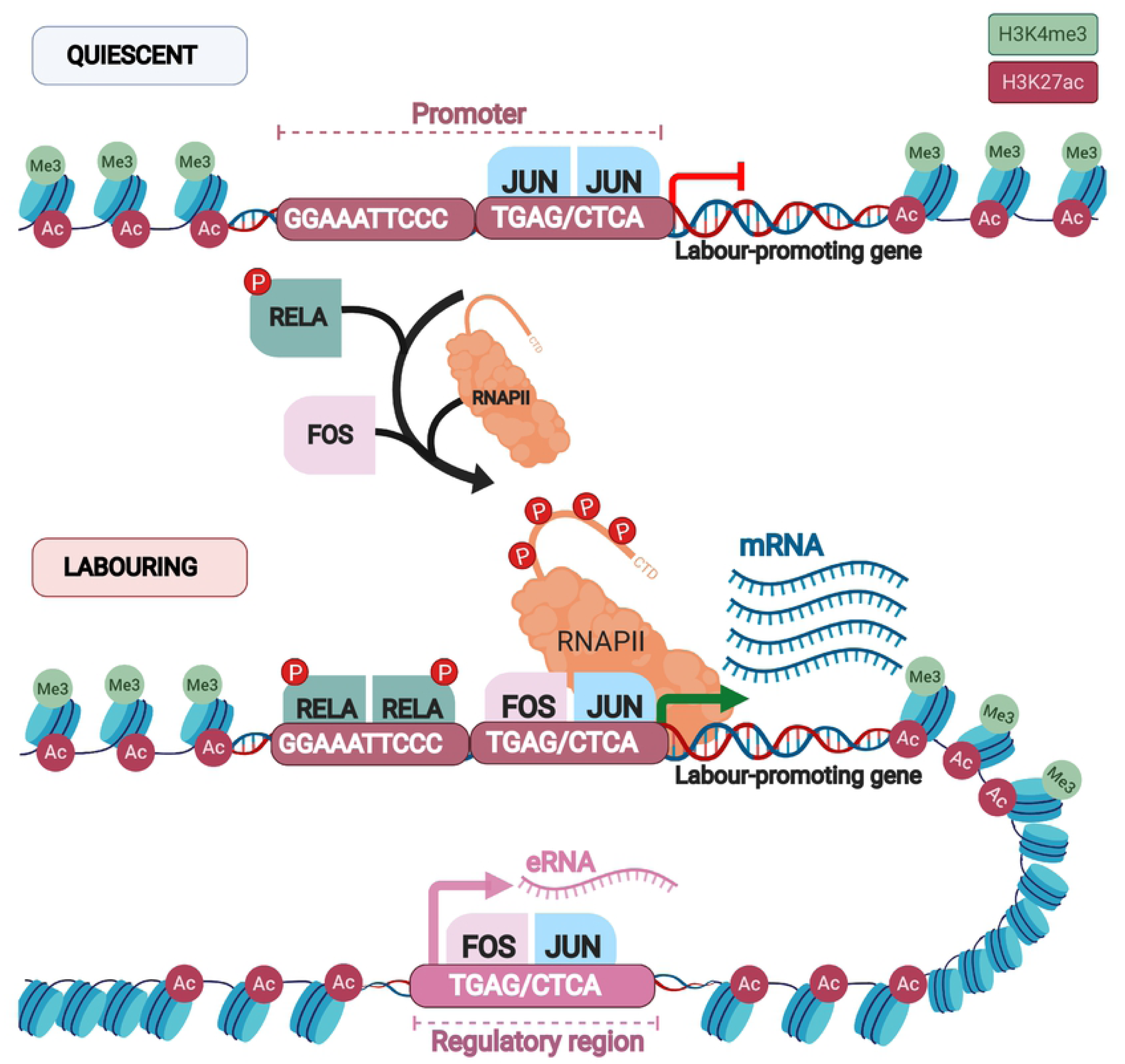
Model of epigenetic priming and transcriptional regulation mechanisms that initiate increased gene expression during labor. Display of the chromatin landscape around typical labor-upregulated genes at quiescent (above) and term laboring (below) stages of gestation. During pregnancy, H3K27ac and H3K4me3 – histone marks typically associated with active genes – are already present at labor-associated gene promoters, thereby priming the epigenome for contractility-promoting transcriptional events in advance of term. However, closer to term, intergenic regions containing AP-1 sequence motifs and modified by H3K27ac enrichment are transcribed, resulting in accumulation of eRNAs. Labor-upregulated gene promoters also contain an enrichment of these motifs, which may allow for binding of homodimerized JUN proteins that are already expressed in the quiescent stage. Conversely, the progression of gestation toward term may result in their replacement by FOS:JUN proteins alongside binding of phosphorylated RELA. We propose that, during labor, these regulatory mechanisms recruit RNAPII to labor-associated gene promoters and enhance transcription through the bodies of these genes.

To our knowledge, this paper is the first to demonstrate that key contractility-promoting genes in the myometrium are up-regulated at term at least in part due to a significant increase in primary transcript abundance. This finding does not preclude the notion that regulation mechanisms act at multiple stages in the expression pathways of labor-associated genes to mediate their expression output. Regulation of *Ptgs2*, for instance, involves miRNA-mediated repressive mechanisms during pregnancy that are halted by reduced expression of miR-199a-3-p and miR-214 as gestation progresses toward term [41,42]. However, our findings reveal that the onset of labor depends on substantial term-restricted transcriptional activity of *Ptgs2*. Whereas our study is the first to investigate these regulatory mechanisms on a genome-wide scale, much of the prior scholarship on labor-associated gene activation events from a single gene perspective has supported this notion. *In vitro* studies have confirmed the regulatory role of select nuclear factor binding sites in critical genes’ promoters: for instance, mutation of an AP-1 factor binding site in the synthetic *Gja1* promoter has been shown to inhibit AP-1 factor-mediated reporter gene expression [21]. Similarly, Khanjani et al. have demonstrated that a 20 base pair-long genomic segment upstream of the human *Oxtr* promoter is required for reporter expression, which is mediated by nuclear factors CCAAT/enhancer-binding protein (CEBP) and RELA [40]. Furthermore, studies’ ChIP-qPCR experiments revealed gestational timepoint-specific binding events that correspond to differential labor-associated gene expression outputs. Renthal et al., for instance, have shown that an intracellular abundance of transcriptional repressors zinc finger E-box binding homeobox proteins ZEB1 and ZEB2 inversely correlates with Oxtr and Gja1 mRNA levels in myometrial cells; furthermore, prominent endogenous binding of ZEB1 and ZEB2 at *Gja1* and *Oxtr* promoters during pregnancy dramatically reduces by term [43]. Finally, increased expression of progesterone receptor A (PRA), a protein critical for contractility in the human myometrial laboring context, is thought to occur due to reduced histone deacetylase 1 (HDAC1) binding and JARID1A histone demethylase enrichment at the PRA promoter [44,45]. These studies suggest that enrichment of activation-prompting histone acetylation and methylation markers at promoters of labor-associated genes guide the transition of the myometrium to a contractile state.

Though these studies were conducted in specific and localized gene contexts, our results regarding the genome-wide enrichment profile of active histone markers over a gestational timecourse were nevertheless surprising. Classical transcription studies that describe the transition of a particular cell type to another state upon subjection to different environmental conditions show evidence of clear histone marker turnover at critical transition-guiding genes [9,46,47]. Contrary to this model, we observed similar trends in H3K27ac and H3K4me3 across all four tested timepoints in mouse myometrial samples, with only subtle acquisition of activating histone marks at significantly up-regulated gene promoters, instead of a clear marker loss or gain according to myometrium state. Furthermore, intergenic regions of labor-associated genes contain H3K27ac peaks that are called as early as day 15, suggesting that not only promoters, but even putative enhancers that can be required for labor initiation may be established during the quiescent phase of pregnancy. Such a molecular set-up can perhaps explain the ease with which labor can occur in advance of term, if portions of the genome that are critical for contractility onset already contain DNA that is open and accessible.

To date, a partial profile of the transcription factors that may bind these open chromatin regions has been put forward. During pregnancy, JUN proteins are known to be present even at early gestational stages [23], prior to the expression of FOS proteins. The interactions of JUN-JUN protein homodimers with co-repressor proteins in quiescent tissues [48] and their potential binding at AP-1 motifs within labor-associated gene promoters may explain why these genes are not activated prior to term. As term approaches, however, FOS sub-family proteins are up-regulated in response to hormonal signals and mechanical stretch stimuli [23,49]. This event results in accumulation of FOS:JUN heterodimers which, we suggest, may bind the same AP-1 motifs in gene promoters, but consequently exert an activating rather than repressive effect on promoters of labor-upregulated genes at this time. Furthermore, the observed enrichment of the RELA motif at these promoters is unsurprising given prior studies highlighting the protein’s role in the pro-labor inflammatory pathway. Increased abundance of phosphorylated RELA immediately prior to the onset of labor [50] suggests that this protein may be a prominent player in the laboring transcription factor network. Furthermore, the other two enriched motifs we found for proteins affiliated with the promotion of gene activation – TCF3 and CTCF – also implicate them in the potential regulation of labor-associated promoter activity. TCF3 may perform a similar coactivator-assisting role that it has been shown to perform in other tissues [36]. CTCF has been proposed to anchor the interactions between gene promoters and distal regulatory elements due to its enrichment at both regulatory regions across cell types [37], a function that this protein may well also enact in the myometrium. Such molecular contributions, as well as the identities of any other transcription factors controlling labor onset, are yet to be determined.

Our study has established a general picture of the chromatin states within the quiescent and contractile myometrial genome and the transcriptional events accompanying the establishment of such states. We have also provided evidentiary support for a broader regulatory role for AP-1 and RELA proteins in regulating these changes across multiple genomic regions. Establishing a more fine-tuned understanding of the molecular basis of birth can allow for a more comprehensive list of therapeutic targets for the prevention of preterm labor in women.

## Materials and Methods

### Animal Model

Bl6 or C57/Bl6 mice used in these experiments were purchased from Harlan Laboratories (http://www.harlan.com/). All mice were housed under specific pathogen-free conditions at the Toronto Centre for Phenogenomics, Canada (TCP) on a 12L:12D cycle, and were administered food and water *ad libitum*. All animal experiments were approved by the TCP Animal Care Committee (AUP# 21-0164-H). Female mice were mated overnight with males and the day on which vaginal plugs were detected was designated as day 1 of gestation. Pregnant mice were maintained until the appropriate gestational timepoint. The average time of delivery was day 19 of gestation. Our criteria for labor were based on delivery of at least one pup from an average number of 14 in two uterine horns.

### Tissue Collection

Animals were euthanized by carbon dioxide inhalation and myometrial samples were collected on gestational day 15, day 19 (term not in labor, TNIL), day 19 during active term (day 19 LAB), and 2-6 hours postpartum (pp). Tissue was collected at 10 a.m. on all days with the exceptions of the labor sample (LAB), which was collected once the animals had delivered at least one pup. The part of uterine horn close to cervix from which the fetus was already expelled was removed and discarded; the remainder was collected for analysis. For each day of gestation, tissue was collected from 4-6 different animals. Mice uteri were placed into ice-cold PBS. Uterine horns were bisected longitudinally and dissected away from both pups and placentas. The decidua basalis was cut away from the myometrial tissue. The decidua parietalis was carefully removed from the myometrial tissue by mechanical scraping on ice, which removed the entire luminal and glandular epithelium and the majority of the uterine stroma. Myometrial tissues were flash-frozen in liquid nitrogen and stored at –80°C. When necessary, myometrial tissues were crushed into fine powder on dry ice prior to subjection to the below listed experimental methods.

### Chromatin immunoprecipitation (ChIP)

Histone marker-targeting ChIP was conducted using the protocol described by Young Lab (younglab.wi.mit.edu/hESRegulation/Young_Protocol.doc), with some necessary modifications for myometrial tissue. Crushed myometrial tissue was fixed in a 1% paraformaldehyde solution at room temperature, a reaction quenched in a 0.125M glycine solution. Cells were rinsed twice with 1X cold PBS and pellets were flash frozen and stored at −80°C until needed. Pellets were washed and samples were lysed in successive lysis buffers, followed by subjection to sonication via the Covaris sonicator with a pulse ON time of 10 s at 30 amps for a total of 30 cycles. Aliquots of sonicated sample were run on a 2% gel to confirm chromatin was sonicated to 300-500 bp size range. Sonicated samples were treated with 10% Triton X-100 and spun down at 4C to pellet debris. Aliquots of cell lysate supernatant to be used as input were stored at -20C.

To bind antibody to magnetic beads, Dynal Protein A and Protein G beads (added in a 1:1 ratio) were washed and resuspended in block solution. Anti-H3K27ac antibody (Abcam, ab4729) or anti-H3K4me3 (Abcam, ab8580) was added, as appropriate, to beads and incubated at 4C with rotation. Beads were again washed and resuspended in block solution. Antibody and magnetic bead mix was added to remaining cell lysate and samples were incubated at 4C overnight with rotation. IP samples were washed at 4C and eluted at 65C. Supernatant was removed from spun-down beads and cross-links were reversed at 65C. Samples were RNaseA-treated at 37C, Proteinase K-treated at 55C, cleaned via phenol-chloroform treatment, and stored in ethanol at -20C overnight. DNA pellets were washed with 80% EtOH and re-suspended in Tris-HCl. Validation of ChIP method was performed using ChIP-qPCR primers (sequences in S11 Table) targeting regions expected to be enriched in our marker of interest.

RNAPII-targeting ChIP was performed as outlined in the supplemental methods in Mitchell and Fraser [51], with modifications for collection of myometrial tissue as performed in case of histone ChIP, and using anti-RNAPII (RPB1 serine 5 phosphorylated form) antibody (Abcam, ab5131).

### ChIP Sequencing and Mapping

ChIP (n=2) and input (n=1) samples from each gestational timepoint – day 15, term-not-in-labor (TNIL), term labor (LAB), and postpartum (pp) – were submitted for single end 50 bp read sequencing using standard Illumina HiSeq 2500 protocols. Reads were quality-checked using FastQC, trimmed with bbduk and mapped to the GRCm38/mm10 mouse reference genome using STAR [52].

### ChIP normalization and peak calling

Peaks were called for each individual replicate using MACS2 broad peak-calling [53]. Significantly conserved peaks in both biological replicates were combined using IDR (Irreproducible Discovery Rate). Differential peak analysis was performed using the diffBind package [54]. Peaks with a foldchange ≥ 4 and adjusted p-value < 0.01 were considered significantly different between day 15 and term labor samples. Peaks with a significant increase in signal intensity in labor samples were linked to the closest gene TSS using bedtools [55]. Normalized ChIP-seq reads (RPM) at promoters (+/-2kb of TSS) of labor-associated upregulated genes in H3K4me3-, H3K27ac-, and RNAPII-targeted samples were quantified using Seqmonk (https://www.bioinformatics.babraham.ac.uk/projects/seqmonk/). Kruskal–Wallis test was used to measure significant (p<0.05) changes in enrichment values (RPM) among different timepoints. Results were plotted using ggplot2 [56]. Sequencing data files were submitted to the Gene Expression Omnibus (GEO) repository (GSE124295).

### Gene expression quantification by RNA extraction and RT-qPCR

Total RNA was extracted from crushed myometrial tissue using Trizol and further DNaseI-treated to remove genomic DNA. RNA was reverse transcribed using the high capacity cDNA synthesis kit (Thermo Fisher Scientific). Target gene expression was monitored by qPCR using exon-intron-boundary-spanning primers (S12 Table) for primary transcript detection, and normalized to levels of total Hist1 mRNA, whose corresponding reference gene was consistently expressed at similar levels across gestational timepoints. Expression levels were calculated against Bl6 or F1 genomic DNA-based standard curve references. All samples were confirmed not to have DNA contamination via non-amplified reverse transcriptase negative samples. Relative expression values were plotted using GraphPad Prism 8. Significant changes in expression were determined by one-way ANOVA with Tukey correction.

### RNA-seq quantification and differential expression analysis

DNAseI-treated total RNA samples isolated from day 15 and term labor mice (n=5 each) were subjected to paired-end sequencing using standard Illumina HiSeq 2500 protocols. Reads were quality-checked using FastQC, trimmed with bbduk and mapped to the GRCm38/mm10 mouse reference genome using STAR [52]. Exon-mapped reads were quantified using featureCounts [57]. Intron reads were quantified using SeqMonk’s active transcription quantitation pipeline (http://www.bioinformatics.babraham.ac.uk/projects/seqmonk/). Alternative transcript counts were summed together for every gene. Intron reads were then imported into DESeq2 [58] for differential expression analysis. Genes with a foldchange ≥ 4 and adjusted p-value < 0.01 were considered significantly changing. Differential RNA expression data was plotted using the EnhancedVolcano package (https://github.com/kevinblighe/EnhancedVolcano). Reads were normalized for gene expression across replicates. Heatmaps were plotted using the pheatmap package (https://cran.r-project.org/web/packages/pheatmap/index.html). Genomic regions of interest for eRNA expression analysis were selected based on intergenic regions featuring H3K27ac peaks with a histone profile displaying a significant increase in signal intensity during labor (foldchange ≥ 4 and adjusted p-value < 0.01). RNA-seq data at these regions were subjected to differential RNA expression analysis by DESeq2. Peaks with fold change ≥ 4 and adjusted p-value < 0.05 were considered as peaks with significantly changing signal intensity from d15 to d19 (labor) and differential eRNA expression was plotted using EnhancedVolcano package.

### Motif enrichment analyses

Enrichment of transcription factor motifs at promoters and intergenic regions of labor-associated genes was performed using HOMER motif analysis tool [35]. Promoter sequences (−1kb of TSS) of upregulated genes at labor were compared against random 1kb input sequences. Intergenic regions containing labor-associated H3K27ac peaks and exhibiting significant upregulation of eRNA expression were compared against random input sequences of varying size. Significantly enriched motifs in the HOMER database were calculated with p-value<0.05.

### Metagene analyses

Genome-wide exon normalized counts were divided into four quartiles according to the average expression of genes (RPM) across replicates in either day 15 or term labor timepoints. Expression quartiles were used to plot the average H3K4me3 and H3K27ac signal and RNAPII coverage at each individual timepoint using ngs.plot [59]. Significantly upregulated and downregulated intron-corresponding reads in term labor samples were used to plot H3K4me3 and H3K27ac signal and RNAPII coverage at all timepoints using ngs.plot.

## Supporting Information

**S1 Fig. Gestational timepoint specific RNA-seq samples cluster based on gestational timepoint of sample collection**. Hierarchical clustering of RNA-seq samples from d15 and d19 when in active labor. Darker colour indicates increased correlation.

**S2 Fig. Proof of ChIP selectivity in myometrial tissues**. Applied anti-H3K27ac ChIP in murine myometrial tissue (target, red) and murine embryonic stem cells (cell control, blue) revealed enrichment or lack of enrichment at select gene targets, as expected. Cell-specific enrichment of this histone mark observed at gene promoters expected to be active predominantly in myometrium rather than embryonic stem cells (left), predominantly in embryonic stem cells rather than myometrium (center), in both cell types (center, right) and in neither cell type (center, left).

**S3 Fig. Enrichment of activating histone marks at gene promoters depends on transcriptional status of genes**. Metagene plots displaying H3K4me3 or H3K27ac enrichment +/-2kb of TSS for genes in expression quartiles reveals increased modification at the promoters of highly expressed genes.

**S4 Fig. Gestational timepoint specific RNAPII ChIP-seq samples cluster based on gestational timepoint of sample collection**. Hierarchical clustering of RNAPII ChIP-seq samples from d15 and d19 when in active labor. Darker colour indicates increased correlation.

**S5 Fig. Enrichment of RNAPII at gene bodies depends on transcriptional status of genes**. Metagene plots displaying RNAPII enrichment at genes in expression quartiles reveals increased association at the promoters and gene bodies of highly expressed genes.

**S6 Fig. Motif enrichment at intergenic H3K27ac regions with increased eRNA during labor**.

**S7 Fig. Motif enrichment at promoters of labor upregulated genes**. Motif enrichment at promoters (1 kb upstream of TSS) of genes with increased expression in labor based on intron reads.

**S1 Table. Genome-wide exon read-based RNA expression values in d15 and term laboring myometrium**.

(XLSX)

**S2 Table. Genome-wide intron read-based RNA expression values in d15 and term laboring myometrium**.

(XLSX)

**S3 Table.** Correlation of gestational timepoint replicates in H3K4me3 and H3K27ac targeted ChIP samples.

| sample | K4me3_D15_A | K4me3_D15_B | K4me3_TNIL_A | K4me3_TNIL_B | K4me3_LAB_A | K4me3_LAB_B | K4me3_1PP_A | K4me3_1PP_B |
| --- | --- | --- | --- | --- | --- | --- | --- | --- |
| K4me3_D15_A | 1 | 0.97 | 0.89 | 0.96 | 0.89 | 0.89 | 0.93 | 0.92 |
| K4me3_D15_B | 0.97 | 1 | 0.9 | 0.97 | 0.9 | 0.91 | 0.94 | 0.94 |
| K4me3_TNIL_A | 0.89 | 0.9 | 1 | 0.91 | 0.9 | 0.85 | 0.9 | 0.85 |
| K4me3_TNIL_B | 0.96 | 0.97 | 0.91 | 1 | 0.91 | 0.91 | 0.94 | 0.94 |
| K4me3_LAB_A | 0.89 | 0.9 | 0.9 | 0.91 | 1 | 0.89 | 0.92 | 0.87 |
| K4me3_LAB_B | 0.89 | 0.91 | 0.85 | 0.91 | 0.89 | 1 | 0.89 | 0.93 |
| K4me3_1PP_A | 0.93 | 0.94 | 0.9 | 0.94 | 0.92 | 0.89 | 1 | 0.91 |
| K4me3_1PP_B | 0.92 | 0.94 | 0.85 | 0.94 | 0.87 | 0.93 | 0.91 | 1 |
| sample | K27ac_D15_A | K27ac_D15_B | K27ac_TNIL_A | K27ac_TNIL_B | K27ac_LAB_A | K27ac_LAB_B | K27ac_1PP_A | K27ac_1PP_B |
| K27ac_D15_A | 1 | 0.86 | 0.83 | 0.85 | 0.83 | 0.8 | 0.79 | 0.73 |
| K27ac_D15_B | 0.86 | 1 | 0.74 | 0.88 | 0.79 | 0.81 | 0.81 | 0.82 |
| K27ac_TNIL_A | 0.83 | 0.74 | 1 | 0.79 | 0.76 | 0.8 | 0.75 | 0.71 |
| K27ac_TNIL_B | 0.85 | 0.88 | 0.79 | 1 | 0.81 | 0.82 | 0.83 | 0.8 |
| K27ac_LAB_A | 0.83 | 0.79 | 0.76 | 0.81 | 1 | 0.85 | 0.87 | 0.79 |
| K27ac_LAB_B | 0.8 | 0.81 | 0.8 | 0.82 | 0.85 | 1 | 0.85 | 0.82 |
| K27ac_1PP_A | 0.79 | 0.81 | 0.75 | 0.83 | 0.87 | 0.85 | 1 | 0.88 |
| K27ac_1PP_B | 0.73 | 0.82 | 0.71 | 0.8 | 0.79 | 0.82 | 0.88 | 1 |

**S4 Table. Reads per million of H3K4me3 at gene promoters**.

(XLSX)

**S5 Table. Reads per million of H3K27ac at gene promoters**.

(XLSX)

**S6 Table. Genome-wide called broad RNAPII peaks in d15, term-not-in-labor, labor, and postpartum myometrium**.

(XLSX)

**S7 Table. Reads per million of RNAPII at gene bodies**.

(XLSX)

**S8 Table. Genome-wide called broad H3K27ac and H3K4me3 peaks in d15, term-not-in-labor, labor, and postpartum myometrium**.

(XLSX)

**S9 Table. H3K27ac peaks displaying a significant increase in read counts in the labor compared to d15 sample**.

**S10 Table. Genome-wide ncRNA expression values in d15 and term laboring myometrium**.

(XLSX)

**S11 Table.**
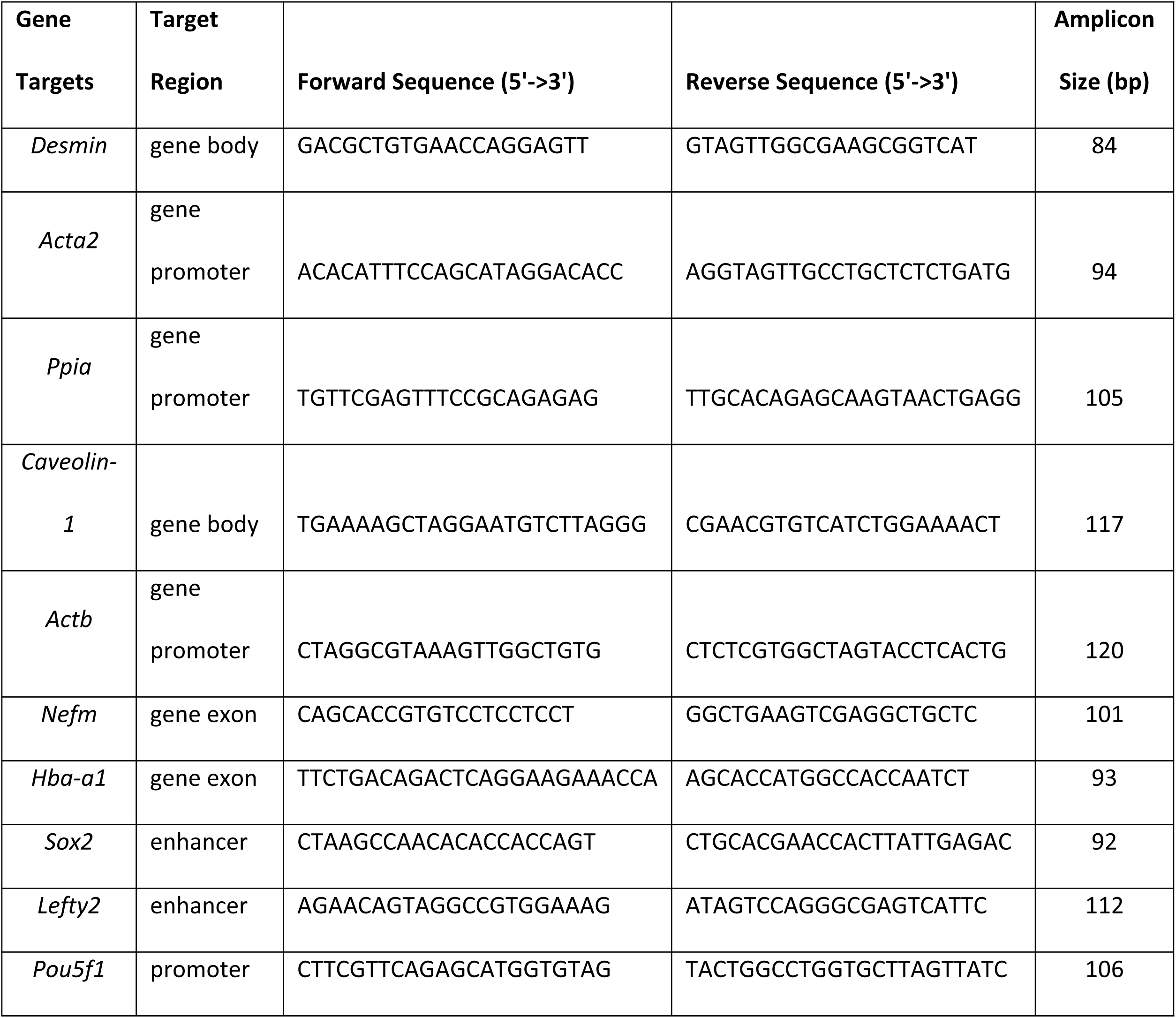
List of primers used in ChIP-qPCR test.

**S12 Table.**
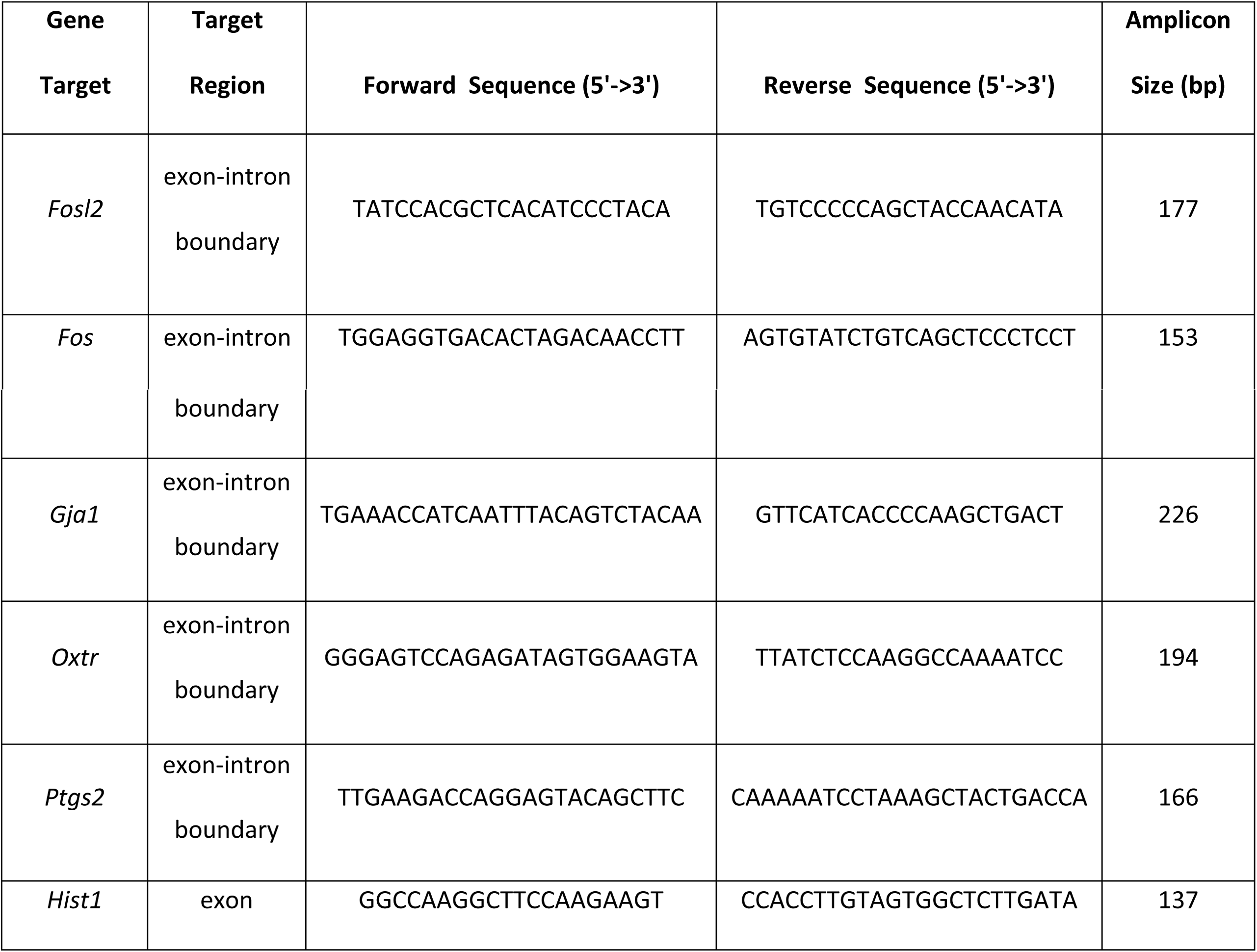
List of primers used in RT-qPCR experiments.

### Acknowledgments

We would like to thank members of the Mitchell, Shynlova, and Wilson labs for their helpful suggestions. This work was supported by the Canadian Institutes of Health Research (FRN 153198, held by J.A.M., O.S., M.W., and V.M.S.), the Canada Foundation for Innovation, and the Ontario Ministry of Research and Innovation (infrastructure grants held by J.A.M.). Studentship funding was provided by the Natural Science and Engineering Research Council of Canada (CGS D held by V.M.S.).

## Author contributions

**Conceptualization:** Jennifer A Mitchell

**Data Curation:** Luis E Abatti, Virlana M Shchuka

**Formal Analysis:** Luis E Abatti, Virlana M Shchuka, Huayun Hou

**Funding Acquisition:** Virlana M Shchuka, Jennifer A Mitchell, Oksana Shynlova, Michael D Wilson

**Investigation:** Virlana M Shchuka, Luis E Abatti

**Methodology:** Jennifer A Mitchell, Oksana Shynlova, Anna Dorogin, Michael D Wilson

**Project Administration:** Jennifer A Mitchell, Virlana M Shchuka, Oksana Shynlova

**Resources:** Oksana Shynlova, Jennifer A Mitchell

**Software:** Luis E Abatti, Huayun Hou

**Supervision:** Jennifer A Mitchell

**Validation:** Virlana M Shchuka, Luis E Abatti

**Visualization:** Luis E Abatti, Jennifer A Mitchell, Virlana M Shchuka

**Writing – Original Draft Preparation:** Virlana M Shchuka, Luis E Abatti, Jennifer A Mitchell

**Writing – Review & Editing:** Virlana M Shchuka, Luis E Abatti, Huayun Hou, Anna Dorogin, Michael D Wilson, Oksana Shynlova, and Jennifer A Mitchell

